# Organ-Founder Stem Cells Mediate Post-Embryonic Neuromast Formation In Medaka

**DOI:** 10.1101/2022.12.16.520711

**Authors:** Karen Gross, Tuğçe Raif, Ali Seleit, Jasmin Onistschenko, Isabel Krämer, Lazaro Centanin

**Affiliations:** Laboratory of Post-Embryonic Lineage Analysis, Center for Organismal Studies, Universität Heidelberg, 69120 Heidelberg, Germany; Heidelberg Biosciences International Graduate School (HBIGS), Universität Heidelberg, 69120 Heidelberg, Germany; Department of Developmental Biology, EMBL Heidelberg, Meyerhofstraße 1, 69117 Heidelberg, Germany

**Keywords:** Post-embryonic Organogenesis, Stem Cell, Neuromast, Lateral Line System, EMT Epithelial-to-Mesenchymal Transition, Medaka

## Abstract

Mammals display a species-specific number, size and location of organs exclusively built during embryogenesis. In fish and amphibians, however, organs must adapt to life-long growth either by expanding in size and/or increasing in number. Here we use neuromasts, small sensory organs that increase in number as fish grow in size, to explore organogenesis during post-embryonic stages. Using iterative imaging, we reveal that post-embryonic organogenesis in the medaka caudal-neuromast-cluster (CNC) is mediated by *organ-founder* stem cells that delaminate from a functional neuromast. *Organ-founder* stem cells undergo epithelial-to-mesenchymal (EMT) transition as shown by molecular markers and cellular rearrangements. Chemokine signaling controls the dynamics of *organ-founder* stem cell delamination, which occurs at a stereotypic position that endures experimental and genetic perturbations. 2-photon laser ablation experiments reveal that organ-founder stem cells are rapidly reconstituted and suggest that these do not constitute a pre-defined population but are rather specified *in situ*. Our findings contribute to better understanding physiological stem-cell mediated organogenesis, a growth strategy present in life-long growing vertebrates. We speculate that a similar strategy could operate in vertebrates with determined-size as a template for pathological conditions like metastasis, where cells detach from their original organ and expand remotely.

## Introduction

Most animals are born with a fixed set of organs that are maintained during their entire life. In these animals, organs have a species-specific number, size and shape, and are specified, formed and enlarged during embryonic development and early post-embryonic life. Fish and some amphibians, however, constitute appealing deviations from this rule. Fish exhibit life-long growth and their organs adapt by expanding in size and/or in number. Within a given species, an older (and longer) fish displays bigger eyes, a bigger brain and more neuromasts than a younger (and shorter) fish. One of the most intriguing questions in the field of post-embryonic organogenesis is whether a post-embryonic organ is formed following the same morphological and molecular steps used during embryonic development. In other words, is there just one cellular and molecular path to generate a given organ? We are interested in the process of post-embryonic organogenesis, i.e how new organs are formed after embryogenesis, with an emphasis on the origin of their founder cells.

Neuromasts are sensory organs distributed along the body of fish and aquatic amphibia (Dijkgraaf, 1963), forming the so-called lateral line (LL) system that detects flow and pressure changes in the surrounding water. Although neuromasts have been used to describe changes in pattern and functional adaptation during post-embryonic stages in zebrafish, medaka and other teleost fish (Nunez et al., 2009; Sapede et al., 2002; Seleit et al., 2021; Wada et al., 2013; Wada et al., 2010), the molecular and cellular basis of post-embryonic organogenesis are just starting to be elucidated. Neuromasts in adult zebrafish and Astyanax are arranged in groups referred to as stitches, which display a dorso-ventral organization along the posterior lateral line (pLL) and are primarily formed by neuromast fission (Ghysen & Dambly-Chaudiere, 2007; Ledent, 2002; Sapede et al., 2002). Pioneer work in early larval stages has shown that post-embryonic neuromasts can also originate from existing neuromast via budding (Wada et al., 2013), for which Wnt signaling induced by the pLL nerve is required. In medaka, post-embryonic neuromast formation can be addressed in the caudal fin where they form the so-called caudal-neuromast-cluster (CNC). Using lineage tracing tools, our group has shown that all neuromasts in the cluster originate from one founder organ that we defined as the P0-neuromast. If the P0-neuromast is ablated during embryogenesis, no neuromast will be found in the CNC in adults (Seleit, Krämer et al., 2017).

Most of our knowledge on the molecular mechanisms to form neuromasts come from studies using the embryonic zebrafish pLL (Aman & Piotrowski, 2008; Chitnis et al., 2012; Ghysen & Dambly-Chaudiere, 2007; Gompel et al., 2001; Haas & Gilmour, 2006; Lopez-Schier et al., 2004; Neelathi et al., 2018; Valentin et al., 2007; Nechiporuk & Raible, 2008; Lecaudey et al., 2008). Briefly, a cluster of cells referred to as pLL-primordium migrates from near the otic vesicle to the caudal fin depositing primary neuromasts along the tail at regular intervals. Primordium migration along the horizontal myoseptum follows a trail of Cxcl12a ligand, which is bound by the receptors Cxcr4b and Cxcr7 expressed by primordium cells (Haas & Gilmour, 2006; Valentin et al., 2007; Donà et al, 2013). The number of embryonic neuromasts deposited is relatively constant in wild type zebrafish, while numerous studies have characterized signaling pathways leading to ectopic embryonic neuromasts when disrupted (Grant et al., 2005; Lopez-Schier & Hudspeth, 2005). Work from our lab has shown that in medaka (*Oryzias latipes*), embryonic neuromasts in the pLL are formed in two sequential morphogenetic waves (Seleit et al., 2017) resulting in *primary* (ventral) and *secondary* (midline) neuromasts distributed in two parallel pLLs. Interestingly, the cytokines *C*xcr4b and *C*xcr7 have been shown to participate in primordium migration and later on, in secondary neuromast formation in medaka (Seleit et al., 2017).

Here we use the formation of the first post-embryonic neuromast in the medaka CNC, the PE1-neuromast (post-embryonic neuromast 1), to characterize cellular and molecular requirements of post-embryonic organogenesis. We show that post-embryonic organogenesis is executed by a few organ-founder neural stem cells that detach from their original organ and migrate to generate a new organ remotely. We link the behavioral modification of organ-founder stem cells with changes in cellular architecture and the expression of epithelial-to-mesenchymal transformation (EMT) marker genes. We also report that chemokine signaling affects the dynamics of organ-founder stem cell delamination, a process that occurs at a stereotypic position. Lastly, we show evidence indicating that organ-founder stem cells are not a pre-defined population but rather a result of local interactions that guarantees a reproducible spatial organization for the post-embryonic organogenesis.

## Results

### Stereotypic formation of a post-embryonic neuromast by *K15+* organ-founder stem cells

Using laser ablation and lineage analysis, we have previously shown that neuromasts in the adult caudal-neuromast-cluster (CNC) originate from *keratin 15* positive (*K15*^*+*^) neuromast stem cells in the P0 neuromast (Figure 1)(Seleit, Krämer et al., 2017). To assess the formation of post-embryonic neuromasts in the CNC dynamically, we performed live-imaging of the P0-neuromast in Tg(*K15*::H2B-RFP) at different stages before, during and after new organ formation (Figure 1). The first morphological change that we detected occurs at the most anterior region of the P0-neuromast (Fig. 1B), where a few *K15*^*+*^ stem cells detach from the founder organ (Fig. 1C,C’, Stage II). The detached cells migrate anteriorly, parallel to and in between the central fin rays (Fig. 1D,D’, Stage III)(N>60 neuromast formation events, N>60 larvae) and eventually generate a new organ just anterior to the P0-founder neuromast, the PE1-neuromast (post-embryonic neuromast 1) (Fig. 1F,F’, Stage V). The process of post-embryonic neuromast formation in the CNC displays variability in timing, but it is extremely stereotypic in terms of directionality of cell migration and position of the post-embryonic organ. Our initial macroscopic analysis suggests that the P0-neuromast contains *K15*^*+*^ stem cells with the capacity to form a new organ. We will refer to those cells as organ-founder stem cells.

**Figure 1.**
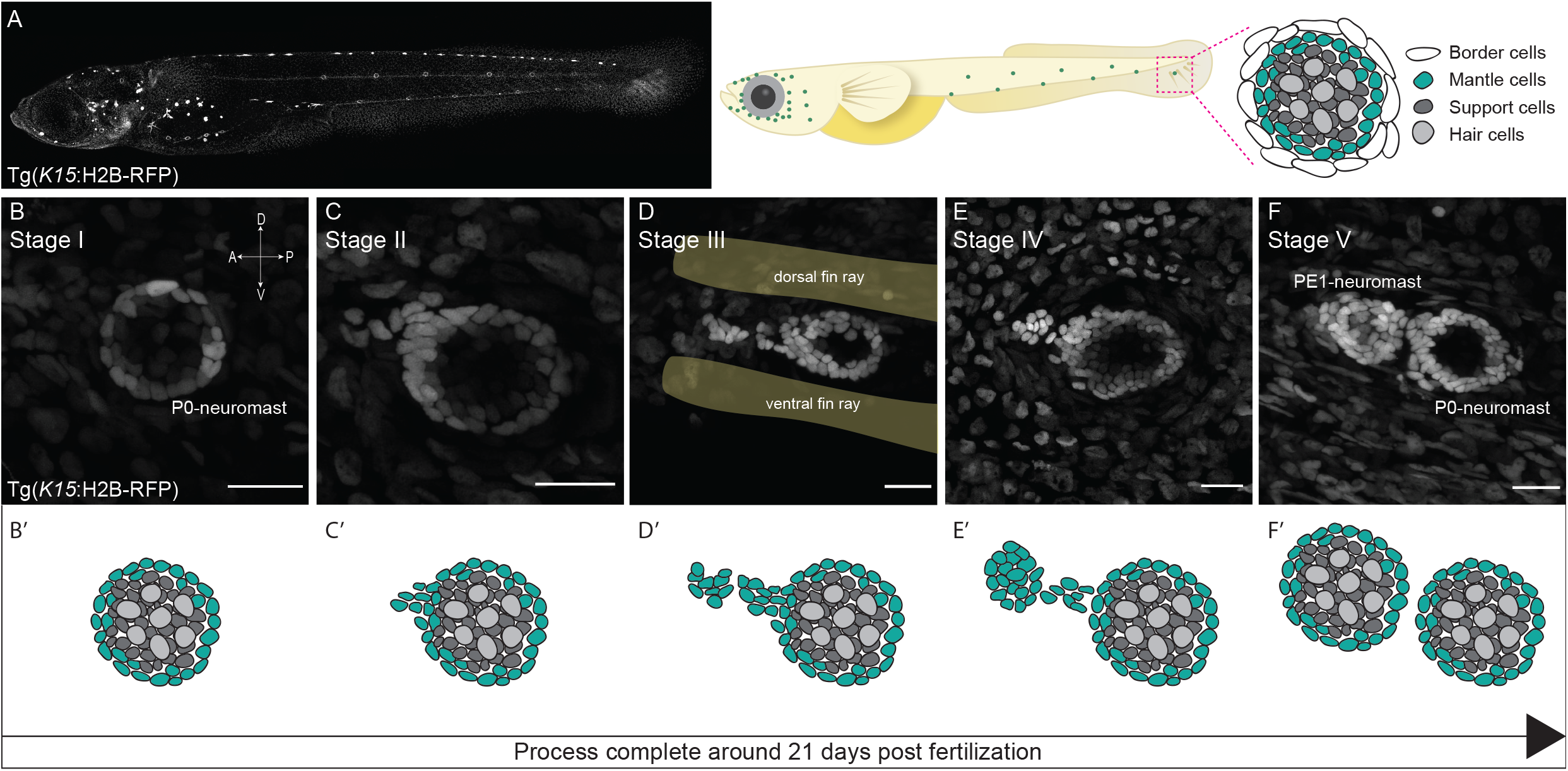
Stereotypic post-embryonic neuromast organogenesis is driven by K15+ organ-founder stem cells. **(A)** Confocal image (left) and scheme (right) of a Tg(*K15*:H2B-RFP) medaka juvenile highlighting neuromasts in the pLL and the P0-neuromast. (**B-F’**) Confocal images (B-F) and schemes (B’-F’) of the P0-neuromast during post-embryonic organogenesis in the medaka CNC. (B-B’) P0-neuromast before the onset of PE1-neuromast formation; notice that the founder organ displays radial symmetry (stage I). (C, C’) Onset of post-embryonic organogenesis is marked by detachment of neuromast stem cells at the anterior border of the P0-neuromast (stage II). (D, D’) Detached stem cells proceed to separate from the P0-neuromast and move in anterior direction, parallel to and in between the central fin rays (yellow shadow) (stage III). (E, E’) Organ-founder stem cells start to coalesce (stage IV). (F, F’) PE1-neuromast formation is completed as revealed by the appearance of differentiated cells (stage V). Scale bars = 20 µm.

### pLL nerve is dispensable for organ-founder stem cell migration and PE1-neuromast formation

We have observed samples in which the formation of PE1-neuromast in the left and in the right pLL happens at different time points after hatch (3/6 fish display different stages of post-embryonic neuromast formation on the left and right side, with a maximum difference of one stage). This variability suggests that organogenesis is not initiated by a systemic signal, but rather triggered by local cues. In zebrafish, it has been reported that new neuromasts originating via budding from previous mature organs require the presence of the lateral line nerve. Innervation is required for the budding process to happen in neuromasts along the pLL, but is not required for other neuromasts like those located on the operculum (Wada et al., 2013; Wada et al., 2010). Since the organ-founder stem cells in the medaka P0-neuromast escape invariably at the anterior region, where the pLL nerve is located (Figure 2A, A’), we hypothesized that the pLL nerve might provide directional cues for cell migration. Therefore, we ablated the pLL nerve connected to the P0-neuromast before the onset of post-embryonic organogenesis using a two-photon laser on double Tg(*Eya1*:mCFP) - to label the pLL nerve - (*K15*:H2B-RFP) - to visualize migrating neuromast stem cells. All ablated fish showed escaping cells from the P0-neuromast (N=7/7 fish), suggesting that the pLL nerve is not necessary for cells to initiate post-embryonic neuromast formation. To exclude that the pLL nerve had already patterned the *escaping point* in the P0-neuromast, we performed laser ablations at earlier stages, before formation of the P0-neuromast (Fig 2B-D). We used the Tg(*Kremen*:mYFP) to label the pLL nerve and the Tg(*Eya1*:EGFP) to assess the differentiation state of the putative newly formed PE1-neuromast. We observed that early pLL nerve ablations did not interfere with the formation of a mature post-embryonic PE1-neuromast that displays post-mitotic neurons as indicated by the EGFP expression from the Tg(*Eya1*:EGFP)(Fig 2C-D)(9/9 fish displayed migrating organ founder stem cells, 7/9 fish generated the PE1-neuromast within two weeks post hatch). Our results indicate that the specification of organ-founder stem cells and the formation of the PE1-neuromasts do not require the presence of the pLL nerve.

**Figure 2.**
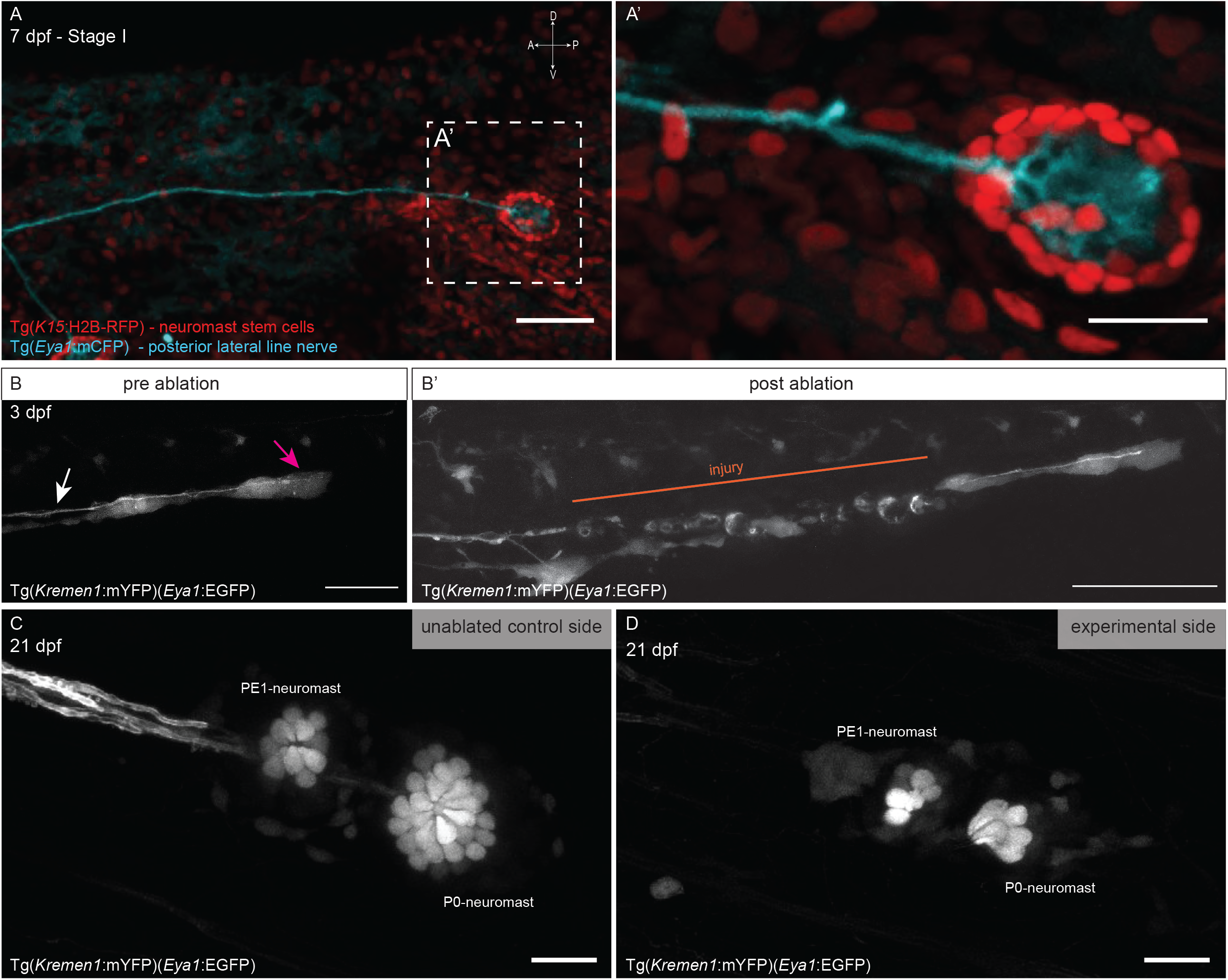
The posterior lateral line nerve is dispensable for PE1-neuromast formation. (**A**) Confocal image of the posterior tail in Stage I double Tg(*Eya1*:mCFP)(*K15*:H2B-RFP) showing the posterior lateral line nerve in cyan and neuromast stem cells in red - detail in A’. Scale bar = 50 µm in A, 20µm in A’. (**B**) Tg(*Kremen1*:mYFP) allows visualization of the pLL nerve (white arrow), while Tg(*Eya1*:EGFP) labels the pLL primordium (magenta arrow) and immature neuromasts at early embryonic stages (3dpf.) Scale bar = 50 µm. (**B’**) Same embryo as B after laser ablation removing parts of the pLL nerve (injury indicated by orange line). Notice the uninjured primordium that continued migration and embryonic neuromast formation. Scale bar = 100 µm. (**C-D**) Tg(*Eya1*:EGFP) labels mature neuromast hair cells in juveniles (20 dpf.) in both PE1 and P0 neuromast on the contra-lateral, non-ablated side (C) as well as on the experimental, ablated side (D). Scale Bar = 20µm.

### Chemokine signaling modulates organ-founder stem cell migration

During the embryonic establishment of the pLL, directional migration of the primordium is controlled by the Cxcr4b/Cxcr7/Cxcl12a signaling pathway. We wondered whether the same pathway that controls primordium migration is re-used by organ-founder stem cells. In situ hybridisation revealed that *cxcl12* mRNA is still present at the onset of post-embryonic organogenesis in the vicinity of the P0-neuromast (Fig.S1). To functionally address whether Cxcr4b is necessary for organ-founder stem cells to migrate away from the P0-neuromast, we generated the *cxcr4b*^*D625*^ mutant via Crispr/Cas directed mutagenesis (Fig.S2). Homozygous *cxcr4b*^*D625*^ larvae display the same phenotype as the previously published *cxcr4b* mutants (Yasuoka et al., 2004), consisting of a disturbed pLL where most organs are not formed due to a non-migratory primordium. The absence of a P0-neuromast in *cxcr4b*^*D625*^ mutants prevented any further analysis on organ founder stem cells.

To mimic the lack of *cxcr4b* in mature organs and in a cell-type specific manner, we used the *K15* promoter to drive expression of the Cxcr7 receptor, a non-signaling (sinking) receptor that competes with Cxcr4b for the Cxcl12 ligand (Aman & Piotrowski, 2008; Dona et al., 2013; Naumann et al., 2010). Although specific for neural stem cells in the mature neuromast, the *K15* promoter drives expression at earlier stages, after primary neuromasts were deposited by the primordium (Seleit et al., 2022). As expected for the over-expression of a functional Cxcr7, Tg(*K15*::Cxcr7) juveniles often lack secondary embryonic neuromasts, which depend on a proper chemokine signaling (Seleit et al., 2017) (Fig.3A,B, green asterisks) (N=24/28 pLLs missing at least one secondary neuromast). We selected fish displaying a phenotype in secondary neuromast pattern and addressed post-embryonic organ formation under reduced Cxcr4b signaling. By following fish via live-imaging at different stages we observed that organ-founder stem cell migration and generation of a PE1-neuromast is delayed in Tg(*K15*::Cxcr7) (Figure 3C-D)(N = 1/6 fish forming a post-embryonic neuromast within 21 dpf (*the WT temporal range)*; N=5/6 fish only reached stage III/IV of post-embryonic organogenesis at 21 dpf). These results suggest that post-embryonic organogenesis in the CNC relies on intact chemokine signaling for its timely completion, similar to primordium migration during embryonic development. Notably, organ-founder stem cells were able to initiate migration in all Tg(*K15*::Cxcr7) juveniles, suggesting that initiation of cell migration is more robust to changes in chemokine signaling than PE1-neuromast organ formation. Later on and as the fish grow, the number of CNC neuromasts increases. Adult Tg(*K15*::Cxcr7) fish display significantly lower CNC neuromast numbers compared to wild type fish (N= 5 WT fish, 3.8 organs per CNC; N= 4 Tg(*K15*:Cxcr7), 2.2 organs per CNC), further suggesting that chemokine signaling via Cxcr4b is indeed involved in post-embryonic neuromast formation in the CNC.

**Figure 3.**
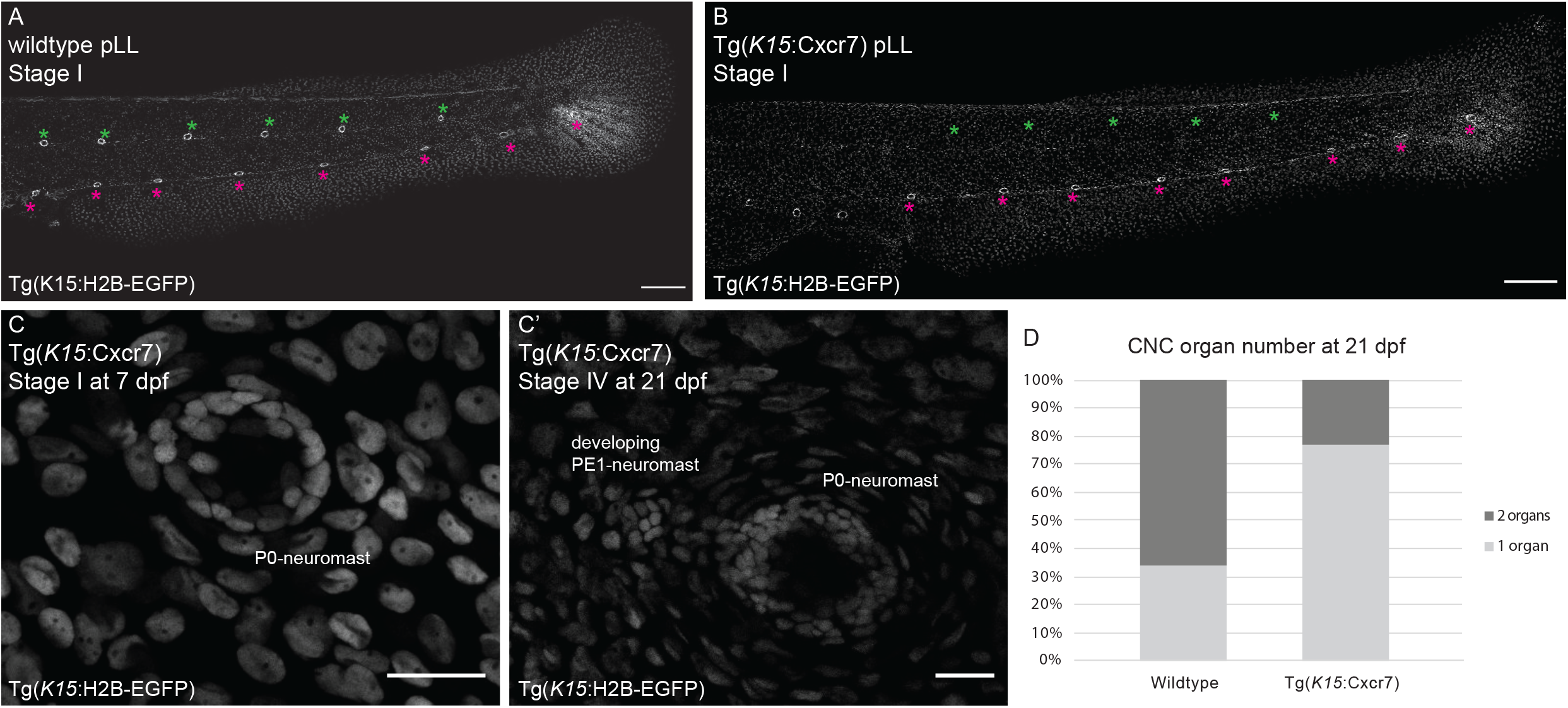
PE1-neuromast formation is delayed in Tg(*K15*:Cxcr7). (**A**) Tg(*K15*:H2B-EGFP) medaka juvenile displays a characteristic zig-zag pattern of neuromasts in the pLL, with secondary neuromasts located along the midline (green asterisks) and primary neuromast located at the ventral side (magenta asterisks). Scale bars = 100 µm. (**B**) Double Tg(*K15*:*Cxcr7*)(*K15*:H2B-EGFP) juveniles are lacking secondary neuromasts along the pLL. Scale bars = 100 µm. (**C**) P0-neuromast of Tg(*K15*:Cxcr7) imaged at 7 dpf (C) and at 21 dpf (C’). Post-embryonic organogenesis has only proceeded to stage IV. Scale bars = 20 µm. (**D**) Quantification of CNC neuromasts of wildtype and Tg(*K15*:*Cxcr7*) fish at 21 dpf.

### Organ-founder stem cells undergo an EMT

During embryonic organogenesis, pro-neuromast cells are deposited by the migrating primordium to form a neuromast. This early step involves the epithelization of neuromast cells, whose first manifestation is the formation of rosettes in the rear end of the primordium (Lecaudey et al., 2008; Nechiporuk & Raible, 2008). Stem cells in the mature neuromast display a characteristic epithelial morphology, with asymmetry along the apical-basal axis (Seleit, Krämer et al., 2017) (Fig.4A, B) (Supplementary Movie1). Our characterization of organ-founder K15+ stem cells in previous sections, indicates that these cells should lose contact with the neighbor stem cells that remain in the P0-neuromast, and suggest that K15+ organ-founder stem cells could be actively migrating away from the P0-neuromast. The transition from a stationary epithelial cell to a migratory cell is known as epithelial-to-mesenchymal transition (EMT), a process where cells lose their apical-basal polarity and acquire migratory properties (Acloque et al., 2009; Lamouille et al., 2014; Moreno-Bueno et al., 2008).

**Figure 4.**
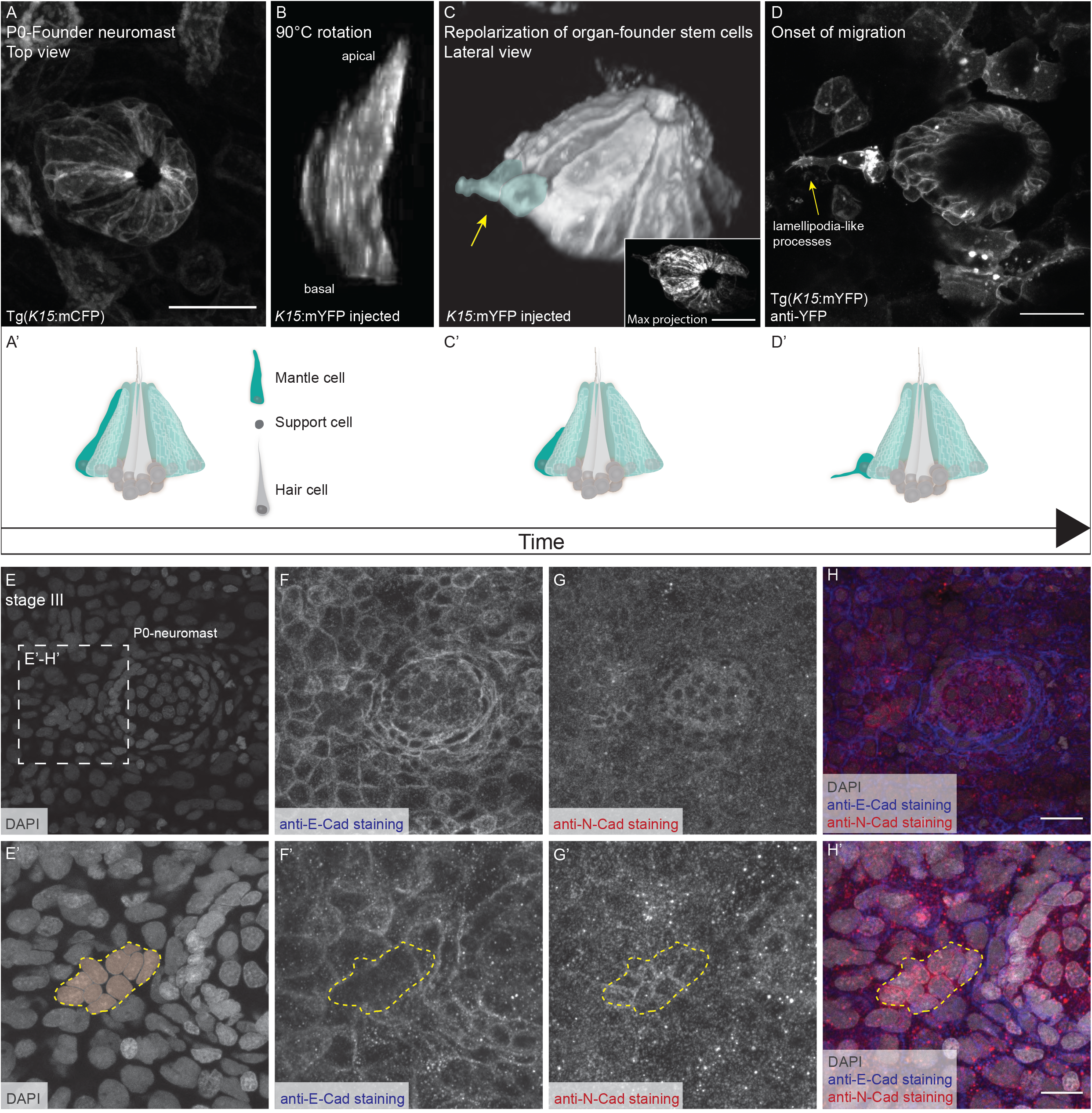
Organ-founder stem cells undergo morphological changes and display an EMT transcriptional signature. (**A**) Top view on P0-neuromast at stage I Tg(*K15*:mCFP) juvenile. All neuromast stem cells are elongated along the apical-basal axis - scheme in A’ from a 90°C angle (side view). (**B**) 3D-reconstruction of two neuromast stem cells in a mosaically labeled fish using *K15*:mYFP highlighting apical-basal elongation - see Supplementary Movie 1. (**C**) 3D reconstruction of a stage II P0-neuromast (side view). Two neuromast stem cells at the anterior border of the P0-neuromast have lost their apical-basal elongation (blue highlight and yellow arrow) defining the onset of Stage II - Schematic depiction in C’. Inlay shows maximum projection of the same P0-neuromast on a top view. Scale bar = 20 µm. (**D**) Organ-founder stem cells form lamellipodia-like processes in the direction of migration (yellow arrow), revealed by immuno-staining against EGFP in Tg(*K15*:mYFP) - schematic illustration in D’. (**E-H**) Immunostainings of EMT related markers and nuclei in P0-neuromast at Stage III; (E-E’) DAPI staining, (F-F’) E-Cadherin, (G-G’) N-Cadherin, (H-H’) Overlay of channels. (E’-H’) Zoom in on migrating organ-founder stem cells. Scale bars = 20 µm.

To address if neuromast organ-founder stem cells indeed undergo an EMT, we used a twofold approach combining morphological and molecular analysis. We examined the morphological changes of organ-founder stem cells during post-embryonic organogenesis using a membrane-tagged fluorescent protein driven by the *K15* promoter in Tg(*K15*:mYFP) (Seleit, Krämer et al., 2017). We observed that at late stage I, K15+ stem cells in the anterior part of the P0-neuromast lose their apical-basal elongation and adopt a mesenchymal-like phenotype (Fig.4C, yellow arrow). Subsequently, the organ-founder stem cells start elongating lamellipodia-like processes in the anterior direction (Fig.4D, yellow arrow), along the path that they will follow during post-embryonic organ formation. In addition and to further characterize the process on a molecular level, we performed immunostaining against E-Cadherin and N-Cadherin, two key indicators of EMT (Fig.4E-H’). Typical hallmarks of EMT are the downregulation of E-Cadherin and the upregulation of N-Cadherin (Kalluri & Weinberg, 2009; Lamouille et al., 2014). We observed that while E-Cadherin is expressed in stationary stem cells of the P0-founder neuromast, it is downregulated specifically in the migrating organ-founder stem cells (Fig.4F, F’). N-Cadherin on the other hand is clearly upregulated in migrating organ-founder stem cells as opposed to stationary neuromast stem cells, in which the signal cannot be detected (Fig.4G, G’). Our morphological and molecular analysis, therefore, indicate that organ-founder stem cells undergo an EMT to migrate away from their initial location, colonize a new position and lead to the formation of a new organ under physiological conditions.

### Invariable exit point for organ-founder stem cells in an ectopic *snail* expression paradigm

Organ-founder neuromast stem cells go through an EMT and start cell migration exclusively at the anterior side of the P0-founder neuromast in medaka. To address whether the system is permissive for ectopic cell migration, we challenged the system aiming at ectopic EMT inductions in the stem cell compartment of the founder organ. The transcription factor Snail1b is one of the key regulators of EMT (Wang et al., 2013); we then generated the Tg(*K15*:snail1b-T2A-H2A-cherry) by using the *snail1b* medaka homologue (Liedtke et al., 2011). In Tg(*K15*:snail1b-T2A-H2A-cherry) juveniles, we observed a disorganized stem cell compartment in pLL neuromasts located along the tail, where stem cells can often be found protruding into the surrounding epithelium (Fig.5A-B’’) (WT: N= 15/35 pLL neuromasts (42.9%); Tg(*K15*:snail1b-T2A-H2A-cherry): N=46/59 (78%) pLL neuromasts displaying SCs protruding from the stem cell compartment). These phenotypes are compatible with a functional *snail1b* and suggest that inducing ectopic EMT can modify the behavior of stem cells in secondary pLL neuromasts. Strikingly, not a single case of ectopic cell migration was observed when we analyzed the P0-neuromast via live-imaging (N= 15 CNCs in 11 fish). In Tg(*K15*:snail1b-T2A-H2A-cherry) fish, cell migration and eventually organogenesis happened exclusively in the anterior direction of the P0-neuromast, like in the *wild type* scenario. However, we noticed a change in the dynamics of post-embryonic neuromast formation driven by Snail1b. While in the WT only 8% of larvae display the post-embryonic PE1-neuromast at 14 dpf (n=12 CNCs; N=12 larvae), that number increases to 33% in Tg(*K15*:snail1b-T2A-H2A-cherry) larvae (n=15 CNCs, N=11 larvae)(Fig. 5C-E). This data suggests that Snail1b alone is not sufficient to promote cell migration at ectopic positions in the P0-neuromast, and indicates that additional information, likely coming from the surrounding environment, is required for organ-founder stem cell migration and post-embryonic organogenesis.

**Figure 5.**
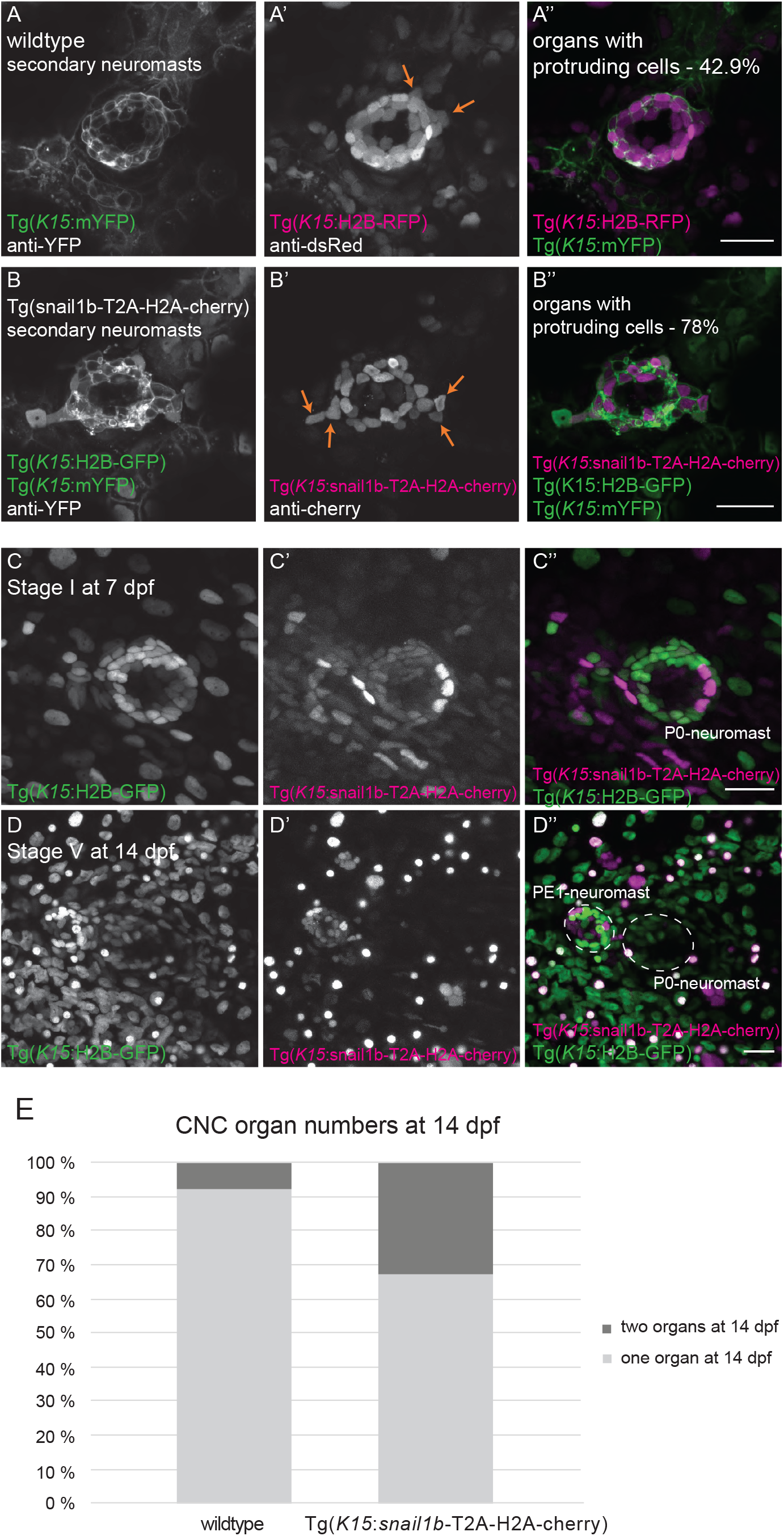
Ectopic expression of Snail1b accelerates post-embryonic organogenesis. (**A-B’’**) Immunostaining against EGFP in Tg(*K15*:mYFP) reveals cellular morphology of neuromast stem cells in secondary neuromasts. (A-A’’) Membrane processes and protruding nuclei (orange arrows) are rarely seen in wild types. (B-B’’) Tg(*K15*:snail1b)display and increased protruding behavior (orange arrows in B’). (**C-D’’**) Live imaging of P0-neuromast in double Tg(*K15*:snail1b-T2A-H2A-cherry)(K15:H2B-EGFP). (C-C’’) Stage I, 7dpf., and (D-D’’) Stage V already at 14 dpf. (**D**) Quantification of organ numbers in the CNC of wildtype and Tg(*K15*:snail1b-T2A-H2A-cherry)(*K15*:H2B-EGFP) at 14 dpf. Scale bars = 20 µm.

### Organ-founder stem cells are reconstituted after ablation

Organ-founder stem cells could be already determined at early embryonic stages, or alternatively, could be generated *in situ* at the anterior boundary of the P0-neuromast. Our results showing that organ-founder stem cell migration cannot be triggered ectopically in the P0-neuromast suggest two possible scenarios. Either the founder organs have a pre-specified reservoir of founder stem cells or alternatively, the local environment creates a permissive corridor only at the anterior position. To test these hypotheses, we aimed at experimentally removing endogenous organ-founder stem cells. To identify the best time point for the ablations, we characterized the proliferative activity of stem cells within the P0-founder neuromast. We used a BrdU pulse of 6 h in samples ranging from the time the P0-neuromast is deposited until the onset of organ-founder stem cell migration (stage III) (5 to 15 dpf) (Fig.6A, A’). This approach revealed a proliferation peak at stage I-II (mean: 6 and 5.4 BrdU+ cells, respectively). Interestingly, we found that most cell divisions occurred at the anterior part of the organ in these stages, constituting the first cellular manifestation of asymmetry along the anterior-posterior axis within the P0-neuromast (Fig.6A’).

**Figure 6.**
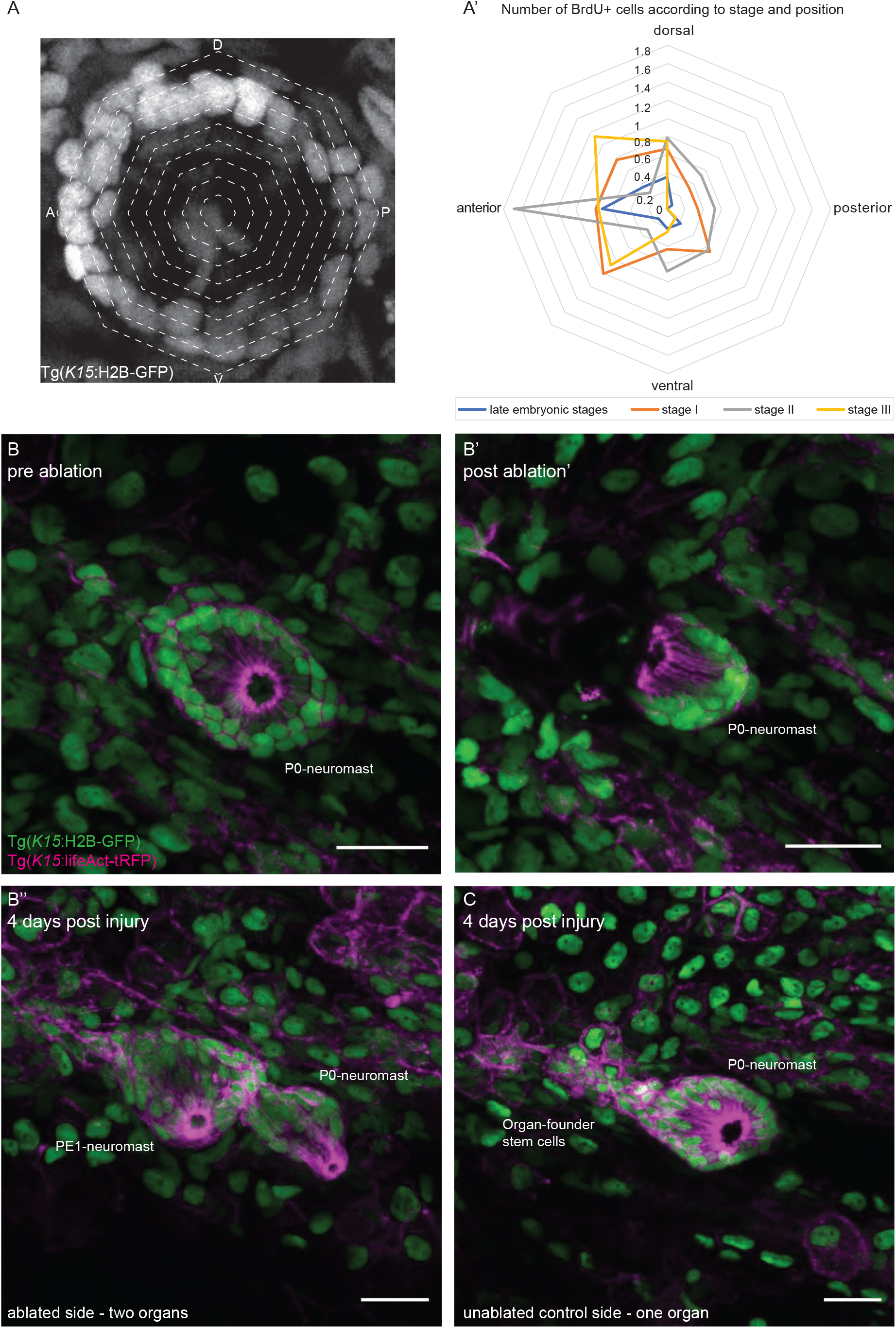
Organ-founder stem cells are reconstituted after laser targeted ablation. (**A**) Analysis of proliferative activity of neuromast stem cells in the P0-neuromast according to stage and position (scheme, left panel). Proliferation peaks at stage I and II, with most BrdU+ cells found in anterior positions (right panel. (**B**) P0-neuromast at early stage II before laser ablation. (**B’**) Same P0-neuromast after ablation of prospective organ-founder stem cells at the anterior half of the organ. (**B’’**) Same fish 4 days post injury, showing already a PE1-neuromast despite ablation of prospective organ-founder stem cells. (**C**) Contra-lateral control side where PE1-neuromast formation has not been completed yet. Scale Bars = 20 µm.

We decided to perform two-photon laser ablations to eliminate neuromast stem cells in the anterior quarter of the P0-founder neuromast at the proliferation peak (Fig.6B-B’). Strikingly, the post-embryonic PE1-neuromast was nevertheless formed from P0-neuromasts in which anterior stem cells were removed (Figure)(N= 7/8 ablated larvae), indicating that the stem cells have been reconstituted. Time-lapse imaging performed immediately after ablations revealed cell rearrangements as the first cellular response (Fig.7). Non-ablated stem cells reorganize, close the gap produced by the ablation and reconstitute organ morphology at the anterior side (Fig. 7C, cyan, blue, green and red dots) while a low number of proliferating K15+ stem cells can be detected at the posterior side (Fig.7C yellow and brown squares) (mean: 0.05 BrdU+ stem cells; N=8 P0-neuromasts). We noticed that in most cases, the ablated P0-founder neuromast was considerably smaller than the non-ablated founder organ in the contralateral side (Fig.6 B’’,C)(N=4/7 larvae). The PE1-neuromast, however, reached the regular size in all cases, regardless of the status of the P0-founder neuromast. Overall, our experiments show that organ-founder stem cells can be reconstituted by other K15+ stem cells, and indicate that the number of stem cells in the P0-founder neuromast is not a critical factor to trigger post-embryonic organogenesis.

**Figure 7.**
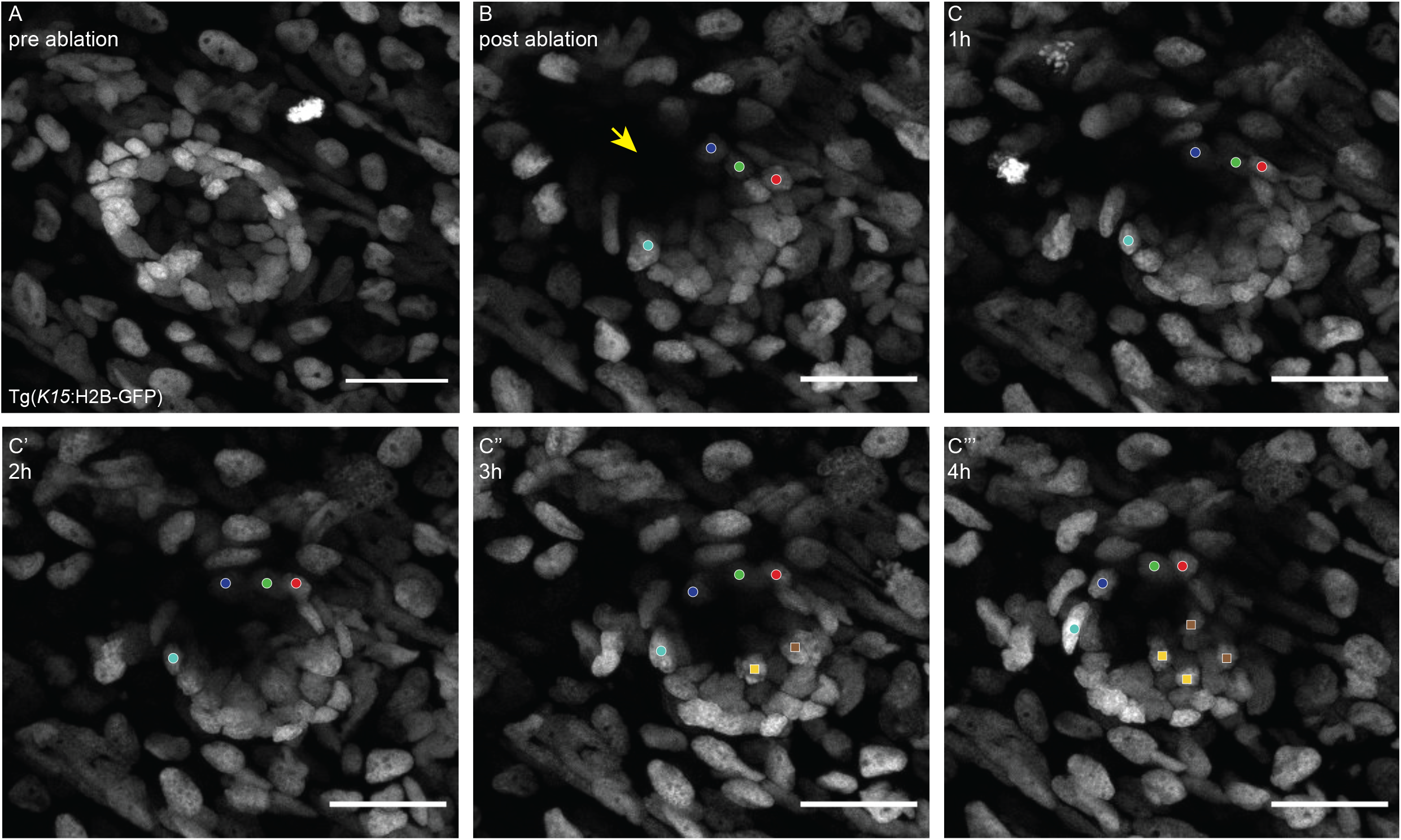
Induction of organ-founder stem cells at anterior position after injury. (**A**) P0-neuromast before ablation of prospective organ-founder stem cells. (**B**) Same organ after laser ablation. (**C-C’’’**) Time-lapse imaging of P0-neuromast after organ-founder stem cell ablation shows stem cell rearrangements to restore organ morphology. Stem cells that were previously positioned dorsal and ventral are now positioned anterior (follow cyan, blue, green and red circles). Stem cell proliferation was seldomly observed (yellow and brown squares) and exclusively occurred at the posterior side of the P0-neuromast - See Supplementary Movie 2.

## Discussion

The work presented here illustrates a fascinating concept: there are multiple ways to build certain vertebrate organs, even within the same organism. Understanding the cellular processes, molecular mechanisms and tissue interactions required for the alternative organogenesis will impact on the current community efforts towards completing organogenesis in culture. In addition to the reports describing different paths to generate neuromasts (Ghysen & Dambly-Chaudiere, 2007; Ledent, 2002; Seleit et al., 2017; Wada et al., 2013), we report here that a group of organ-founder stem cells detaches from its current environment via an epithelial-to-mesenchymal transformation and travels away to generate a new organ remotely. This physiological process resembles molecular steps observed during pathological metastases in mammals. If, as previously proposed (Shubin, 2008), the an-amniote body plan and physiology are the foundation of the amniote body, it is plausible that ancient physiological processes that were in place to cope with a permanently growing body are still present, albeit suppressed, and can be hijacked during pathological conditions in organisms with a determined size like humans.

### Metastatic-like organogenesis vs. neuromast budding

Post-embryonic neuromast formation by *budding* has been described before in amphibia and some teleost fish species (Stone, 1937; Wada et al., 2013). The process we are describing here is mechanistically distinct from neuromast budding in the zebrafish caudal fin, which crucially depends on innervation (Wada et al., 2013). There, the axons innervating budding neuromasts express the Wnt activator R-Spondin, resulting in neuromast cell proliferation and bud formation. We show that in contrast to that, innervation is dispensable for initiation of accessory organogenesis in the medaka CNC. Even in the absence of the pLL nerve, organ-founder stem cells in medaka migrate away from the founder organ and form the PE1-neuromast. Whether neuromast stem cell proliferation in the accessory organ in medaka is also driven by Wnt signaling still needs to be examined, as well as the putative source of the proliferative signal. Another important difference is the PE1-neuromast does not originate as a bud from the P0-neuromast, but rather it is generated from K15+ neural stem cells that migrate away from the founder organ. We believe that this cellular mechanism resembles the metastatic activity of tumor cells in mammals, and hence, would name it *metastatic-like* organogenesis.

### Chemokine signaling impacts on the temporal dynamics of PE1-neuromast formation

Our results using cell type-specific overexpression of Cxcr7 in neuromast stem cells suggest that chemokine signaling is indeed participating during post-embryonic organogenesis in the medaka CNC. Tg(*K15*:Cxcr7) juveniles display a substantial delay in PE1-neuromast formation, although eventually these will be formed at their proper position. The *delayed* phenotype could be explained by Cxcr7 affecting either a) initiation, or b) directionality, of organ-founder stem cell migration. If the chemokine signaling impacts on the initiation of organ-founder stem cell migration, the competition between Cxcr7 and Cxcr4b for the same ligand would result in a less efficient process. It might take more time for the organ-founder stem cells to accumulate the molecular effectors to successfully engage in migration. Our observations comparing wild type P0-neuromasts within the same fish indicate indeed that the migration-initiation phase displays temporal variability compatible with what we observed in the Tg(*K15*:Cxcr7). On the other hand, chemokine signaling might not affect migratory behavior itself, but rather directional movement. During the establishment of the embryonic lateral line, *cxcr4b* mutants exhibit primordium migration defects. These defects are not due to failure of initiation of migration, but to a loss of directionality (Haas & Gilmour, 2006). In our system, the distance traveled by organ-founder stem cells is rather short, which might enable organ-founder stem cells to eventually migrate in the right direction by a permissive corridor built by the surrounding tissue.

### Plasticity of organ-founder stem cell fate

Out of all the cells in the CNC founder neuromast only a subset of stem cells located at a stereotypic position generate an accessory organ. The two main possible scenarios are: i) some stem cells are pre-specified to become organ-founder stem cells, and ii) all stem cells have the same capacity to generate a new organ, but the system is only permissive in anterior positions. We tackled the problem of pre-specification vs. plasticity by ablating prospective organ-founder stem cells and assessing the impact on organogenesis. Upon ablation of prospective organ-founder stem cells (stem cells in the anterior part of the organ), other neuromast K15+ stem cells compensated for their loss and acquired an organ-founder stem cell fate. The reconstitution of a stem cell population after injury or ablation has indeed been observed in other mammalian stem cell systems like the intestine (Tian et al., 2011) and the hair follicle (Rompolas et al., 2013) among others. Reconstitution of the system happens by recruiting a reserve stem cell population (Tian et al., 2011) or via other cells in the lineage that locate into the empty niche and acquire stem cell properties (Rompolas et al., 2013; Tetteh et al., 2016). Here we see that other stem cells within the same organ reconstitute the organ-founder function by cellular rearrangements.

An interesting observation after the ablation experiments is that in 50% of cases, the P0-neuromast is smaller than the newly generated PE1-neuromast, a situation that we never observe in the un-injured case. Neuromasts display a well reported capacity for regeneration, even after losing most of their cells (Pinto-Texeira et al, 2015; Seleit, Krämer et al, 2017; Viader-Llargués et al, 2018). Our results indicate that recruiting stem cells to generate a new organ outweighs the drive to immediately reconstitute missing cell numbers. This again reinforces the notion that new organ-founder stem cells do not originate from proliferation of remaining cells, but rather that remaining cells directly acquire an organ-founder stem cell fate.

### Tissue interactions induce stem cell heterogeneity

The microenvironment has been shown in different model systems to be instructive for cell behavior, such as quiescence, proliferation, differentiation and migration via both mechanical and biochemical signals (Choi et al., 2018; de Lucas et al., 2018; Fiore et al., 2018). Our observations show that the onset of post-embryonic organogenesis is marked by proliferation, cellular rearrangement and migration of neuromast stem cells exclusively at the anterior part of the P0-neuromast. Interestingly, our aims at buffering the molecular asymmetry within the P0-neuromast by ectopic expression of *snail1b* in neuromast stem cells did not abolish asymmetric stem cell migration and post-embryonic organogenesis. Instead, stem cell migration and post-embryonic organogenesis occurred exclusively anterior, albeit slightly faster than in the wildtype. This result indicates that only the microenvironment surrounding the P0-neuromast at the anterior border is permissive for cell migration and eventually post-embryonic organogenesis. We have noticed that in Tg(*K15*:snail1b) larvae epithelial cells surrounding the P0-neuromast display nuclear condensation (Fig. 5D-D’’) suggesting cell death. This in turn might have non-autonomous effects on organ-founder stem migration by changing the architecture of the surrounding tissue. Our stem cell ablation experiments demonstrate that the anterior microenvironment is not only permissive for cell migration, but also stimulates heterogeneity among neuromast stem cells by inducing organ-founder stem cell fate specifically in the anterior region. In addition, the reconstitution of the organ-founder stem cell pool allows us to state these are not a pre-specified population within K15+ neuromast stem cells, but are rather induced *in situ*. The functional output of this rationale is an adaptable system able to react dynamically to the local needs of the growing tissue, inducing the formation of organ-founder stem cells to give rise to a new organ in the vicinity with certain autonomy from the rest of the neuromasts in the organism.

## Material and Methods

### Fish maintenance

Medaka fish (*Oryzias latipes*) used in this study were kept according to local animal welfare standards (Tierschutzgesetz §11, Abs. 1, Nr. 1) as closed stocks and maintained in a recirculating system at 28°C on a 14 h light/10 h dark cycle (Tierschutzgesetz 111, Abs. 1, Nr. 1, Haltungserlaubnis AZ35–9185.64 and AZ35–9185.64/BH KIT). Animal experiments were conducted according to European Union animal welfare guidelines. The published transgenic lines used in this study are : Tg(*Eya1*:mCFP), Tg(*Eya1*:EGFP) (Seleit et al, 2017), Tg(*K15*:H2B-RFP), Tg(*K15*:H2B-EGFP) (Seleit, Krämer et al, 2017), Tg(*K15*:LifeAct-tagRFP) (Seleit et al, 2022). Tg(*Kremen1*:mYFP) was generously provided by J. Wittbrodt’s lab.

### Generation of Tg(*K15*:*snail1b*-T2A-H2Acherry)

The plasmid I-SceI/*K15*:snail1b-T2A-H2Acherry/I-SceI was generated by amplifying the *snail1b* CDS and H2Acherry separately with specific primers containing additional overlapping sequences to allowed for a further fusion PCR : GTCGACGTCTTCCAACATTTACGCACG (*snail1b* forward with SalI overhang), CCTCCACGTCACCGCATGTTAGAAGACTTCCTCTGCCCTCCGCTGAGGCACAGCAGGA (snail1b CDS reverse with partial T2A site overhang), TCTTCTAACATGCGGTGACGTGGAGGAGAATCCCGGCCCTATGGCAGGTGGAAAAGCAGGT, (H2Acherry forward with partial T2A site overhang) and GCGGCCGCTGCATTCTAGTTGTGGTTTGTCCA (H2A-cherry reverse with NotI overhang).

The fusion construct was cloned under the *K15* promoter in a I-SceI containing vector. The final construct was injected into 1-cell stage medaka embryos using meganuclease mediated integration and screened for expression of H2A-Cherry in epithelium and neuromast cells to generate Tg(*K15*:snail1b-T2A-H2Acherry).

### Generation of *cxcr4b*^*D625*^ mutant line

The *cxcr4b*^*D625*^ mutant line was generated using two gRNAs (GUGAAAACCUGGUACUUCGGAGGGTGAAAACCTGGTACTTCGGAGG, CAAGUGGAUUUCUAUCACCGAGGCAAGTGGATTTCTATCACCGAGG) targeting the coding sequence of *cxcr4b*, resulting in a deletion of 625 base pairs. Design of gRNAs was done using CCTop (Stemmer et al., 2015). Cloning of gRNAs was done as described before (Seleit et al., 2022; Seleit, Krämer et al., 2017; Stemmer et al., 2015) and used at a concentration of 15 ng/µl together with 150 ng/µl Cas9 mRNA. We injected 1-cell stage medaka embryos from Tg(*Eya1*:EGFP) or Tg(*K15*:lifeAct-tRFP), screened the resulting larvae for lateral line phenotypes and grew them to adulthood. We identified carriers by genotyping the offspring using the following oligos: CCCCTCCATTGTTCTTGTCGC and CCCAGATTTTCCACGCAGACT. The mutant line was established and subsequently maintained via outcrosses to wild type fish.

### Generation of Tg(*K15*:*Cxcr7*)

The I-SceI/*K15*:Cxcr7/I-SceI construct was generated by cloning the full-length coding sequence of Cxcr7 in a I-SceI containing plasmid (Thermes et al., 2002) downstream of the *K15* promoter. We generated the Tg(*K15*:Cxcr7)(*K15*:H2B-GFP) by injecting 1-cell stage medaka embryos together with an additional plasmid containing a I-SceI/*K15*:H2B-GFP/I-SceI and the I-SceI enzyme.

### Live-imaging

Live fish were tranquilized in Tricaine (MS-222) (Sigma-Aldrich) for about 5-10 min before being mounted in 0.6% low-melting agarose (Roth) in glass bottom dishes (MatTek). Imaging was performed using confocal laser-scanning microscopy (Leica SP5 II and Leica TCS SPE). Image analysis was performed using FIJI (Schindelin et al., 2012).

### Multi-photon laser ablation

For targeted laser ablation the fish were mounted as described above. Multi-photon laser ablation was performed using the Leica SP5. The multi-photon laser was used with the option ‘Area ablations’ and the 880 nm wavelength at 65% laser power for 8-10 seconds at a region of interest. Successful ablations were assessed immediately after.

### Immunohistochemistry

Immunohistochemistry was performed as described before (Centanin et al., 2014). Primary antibodies used were rabbit-**α**-GFP (Invitrogen, A11122, 1/500), rabbit-∝ -mCherry (abcam, ab167453, 1/500), mouse-**α**-E-Cadherin (BD Biosciences, 610181, 1/500), rabbit-**α**-N-Cadherin (abcam, ab76011, 1/250). Secondary antibodies used were goat-**α**-rabbit-488 (Life Technologies, A-11034, 1/500), donkey-α-mouse-647 (Jackson/Dianova, 715-605-151, 1/500), goat-**α**-rabbit-549 (Jackson, 112-505-144, 1/500), goat-α-rabbit-647 (Invitrogen, A21245, 1/500). In order to label all nuclei, the samples were incubated with DAPI (1/500).

### BrdU incorporation and staining

Fish at different stages were incubated with 2.5 mM BrdU (Sigma-Aldrich) for 6 hours at room temperature and fixed immediately after. Samples were co-stained with DAPI (1/500), typically prior to denaturation of the DNA required for the BrdU staining protocol. We included an additional antigen retrieval step before the blocking step; antibodies used were rat-α-BrdU (Abcam, ab6326, 1/200) and goat-α-rat-488 (Jackson, 112-485-143, 1/500).

## Acknowledgements

We would like to thank past and current members of the Centanin lab, S.Lemke, J-R Martinez Morales for critical input on the manuscript and for fruitful discussions; J. Wittbrodt for the generous access to equipment, J. Wittbrodt’s Lab for sharing the unpublished Tg(*Kremen1*:mYFP), NBRP Medaka (https://shigen.nig.ac.jp/medaka/) for materials, C. Loosli for technical assistance with genotyping, and E. Leist, A. Sarraceno and M. Majewski for assistance regarding fish maintenance.

## Figure legends

**Figure S1 - *cxcl12a* is present at the onset of post-embryonic organogenesis**. (**A**) Tail of a wildtype juvenile fish at the onset of post-embryonic organogenesis. *cxcl12a* mRNA can be detected along the myoseptum and in the caudal fin. (**B**) Zoom-in showing that *cxcl12a* mRNA can be detected in the vicinity of the P0-neuromast. Scale Bar = 100 µm.

**Figure S2 - *cxcr4b***^***D625***^ **mutants contain long deletion and display severe pLL phenotype**.

(**A**) Schematic depiction of *cxcr4b* wildtype locus and *cxcr4b*^*D625*^ mutant allele with position of guide RNAs and genotyping primers. (**A’**) Sequencing read of *cxcr4b*^*D625*^ mutant allele. (**B-C**) Tg(Eya1:EGFP) medaka juvenile - neuromasts are highlighted in cyan, complete EGFP+ pattern displayed in green. Wildtype pLL displays alternating primary and secondary neuromasts along the tail (B) while in *cxcr4b*^*D625*^ mutants the pLL is severely truncated (C, white arrow). Notice that the P0-neuromast is not formed in *cxcr4b*^*D625*^ mutants.

**Supplementary Movie 1 - Neuromast stem cells are elongated along the apical-basal axis**. 3D reconstruction of a subset of neuromast stem cells that were labeled by mosaic expression of a *K15*:mYFP plasmid.

**Supplementary Movie 2 - Re-arrangement of organ-founder stem cells to the anterior position after ablation**.

Live imaging of a Tg(*K15*:H2B-EGFP) juvenile immediately after laser ablation of the anterior stem cells on a late Stage I P0-neuromast.

